# Notch signaling is required for survival of the germline stem cell lineage in testes of *Drosophila melanogaster*

**DOI:** 10.1101/682773

**Authors:** Chun L. Ng, Qian Yue, Schulz Cordula

**Affiliations:** University of Texas Southwestern Medical Center, Dallas, TX 75390; University of North Georgia, Department of Biology, Oakwood, GA 30566; University of Georgia, Department of Cellular Biology, Athens, GA 30602

## Abstract

In all metazoan species, sperm is produced from germline stem cells. These self-renew and produce daughter cells that amplify and differentiate dependent on interactions with somatic support cells. In the male gonad of *Drosophila melanogaster*, the germline and somatic cyst cells co-differentiate as cysts, an arrangement in which the germline is completely enclosed by cytoplasmic extensions from the cyst cells. Notch is a developmentally relevant receptor in a pathway requiring immediate proximity with the signal sending cell. Here, we show that Notch is expressed in the cyst cells of *wild-type* testes. Notch becomes activated in the transition zone, an apical area of the testes in which the cyst cells express stage-specific transcription factors and the enclosed germline finalizes transit-amplifying divisions. Reducing the ligand Delta from the germline cells via RNA-Interference or reducing the receptor Notch from the cyst cells via CRISPR resulted in cell death concomitant with loss of germline cells from the transition zone. This shows that Notch signaling is essential for the survival of the germline stem cell lineage.

## Introduction

The Notch signaling pathway is highly conserved and plays versatile roles in development, such as nervous system formation, cardiac patterning, and sensory hair formation (Guruharsha et al., 2012; Jain et al., 2010; MacGrogan et al., 2010). In many developmental contexts Notch specifies cell fate decisions. In the developing vertebrate eye, for example, Notch regulates which cells develop into glial cells and which develop into optic neurons (Dorsky et al., 1997; Wang et al., 1998). During *C. elegans* vulva development, Notch prevents nearby cells from becoming central vulval cells (Berset et al., 2001). In other developmental contexts Notch regulates the survival of cells. In the murine nervous system, loss of Notch results in the death of progenitor cells and newly differentiated cells (Mason et al., 2006). Notch signaling has also been associated with cell survival in B-cell malignancies, prostate cancer cells, and myeloma cells (Nefedova et al., 2008; Ye et al., 2012; Zweidler-McKay et al., 2005).

The canonical Notch signaling pathway is rather simple. Upon activation, the Notch receptor is proteolytically cleaved causing the release of the intra-cellular portion of the protein, called the Notch intra-cellular domain (NICD). The NICD enters the nucleus and joins a protein complex bound to chromatin altering the transcription of target genes. This complex includes the transcription factors Suppressor of Hairless (Su(H)) in *Drosophila melanogaster* and Mastermind, as well as other potential co-regulators (Figure 1A). Additional levels of regulation are added to the pathway via receptor-ligand internalization, post-translational modification, and protein stability (Bray, 2016; Hori et al., 2013).

**Fig. 1:**
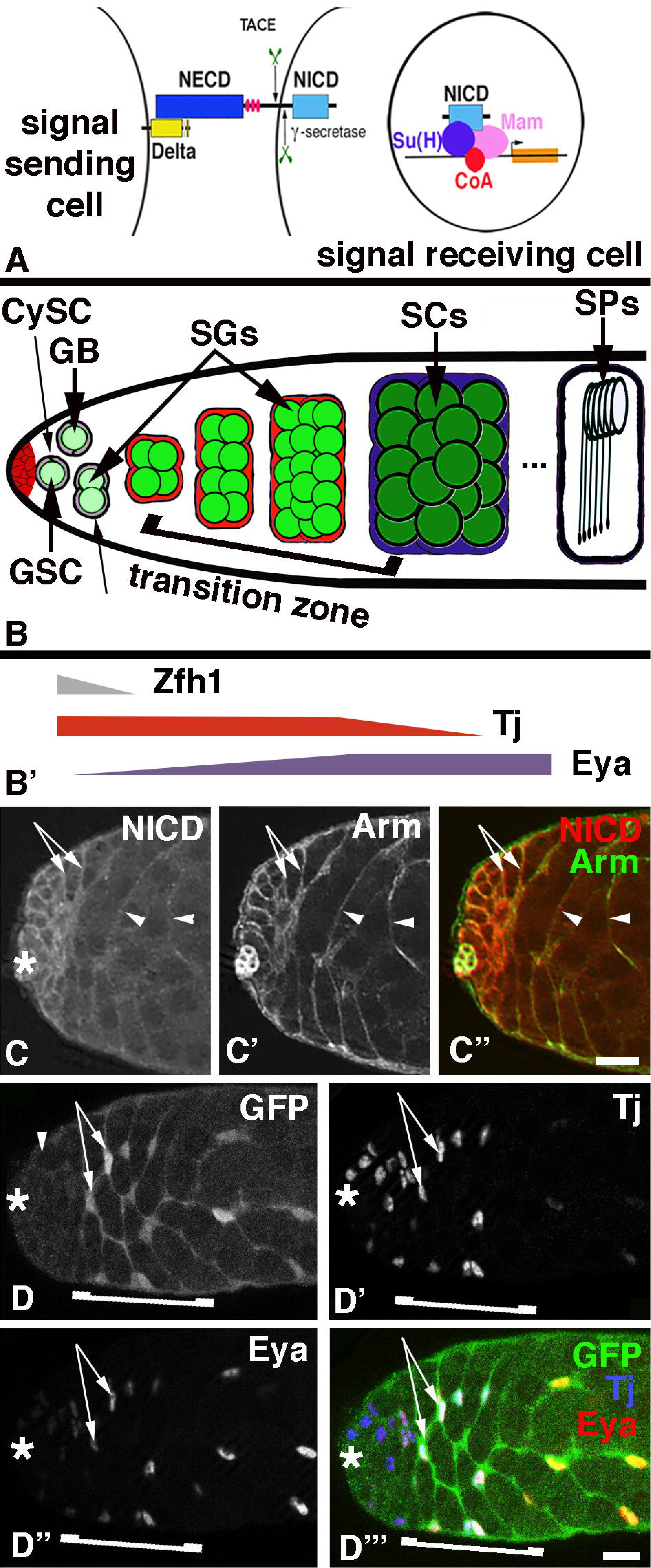
Notch signaling is activated in the transition zone. A) Cartoon depicting the canonical Notch signaling pathway. NECD: Notch extra-cellular domain, NICD: Notch intra-cellular domain, Su(H): Suppressor of Hairless, Mam: Mastermind, CoA: transcriptional co-regulator, TACE and γ-secretase: proteases cleaving the Notch receptor. B) Cartoon of spermatogenesis with a focus on the apical region. GSC: germline stem cell, GB: gonialblast; SGs: spermatogonia; SCs: spermatocytes; SPs: spermatids; CySC: Cyst Stem Cell, CCs: cyst cells, bracket: transition zone. B’) Arrows represent the regions of the testis in which indicated transcription factors are expressed in the CySC and cyst cells. Color-coding corresponds to the colors in B). C-D’’’) Asterisks mark the apical tips of the testes, scale bars: 30 μm. C-C‘’) Apical region of a *wt* testis showing C) NICD, C’) Arm, and C’’) their co-localization on the cyst cell membranes, as indicated by arrows and arrowheads. D-D’’’) The D) NRE-eGFP reporter for Notch activation co-localized with D’) Tj and D’’) Eya within the cyst cell nuclei of a *wt* testis (D’’’). The transition zone is depicted by a bracket.

While vertebrates have several Notch receptors and ligands, the *Drosophila* genome only contains one Notch receptor and two ligands, Delta (Dl) and Serrate. Both, the receptor and the ligands are transmembrane proteins (Kopan and Ilagan, 2009). Thus, in order for Notch signaling to occur the ligand-expressing cells have to be in intimate contact with the receptor-expressing cells. Such a cellular architecture is found in spermatogenesis where germline cells are associated with and surrounded by somatic support cells (Hardy et al., 1979; Griswold, 1998). *Drosophila* testes contain Germline Stem Cells (GSCs) that produce sperm cells and Cyst Stem Cells (CySCs) that produce somatic support cells, the cyst cells. Both stem cell populations are arranged around a group of somatic hub cells at the apical tip of the testis (Figure 1B). GSCs and CySCs undergo constant cell divisions that result in asymmetric outcomes, the stem cell daughters that remain attached to the hub cells become new stem cells, while the stem cell daughters that are displaced away from the hub become gonialblasts and cyst cells, respectively (Hardy et al., 1979; Yamashita et al., 2003). One gonialblast and two cyst cells form a cyst. During this process, the cyst cells grow cytoplasmic extensions around the gonialblast to fully enclose it (Sarkar et al., 2007; Schulz et al., 2002). The two cyst cells serve as an adhesion and signaling center for the developing germline cells. Accordingly, defects in cyst formation result in abnormal proliferation and differentiation of the germline and cyst cells (Fairchild et al., 2017; Kiger et al., 2000; Matunis, 1997; Sarkar et al., 2007; Schulz et al., 2002; Tran et al., 2000).

During subsequent cyst differentiation, the gonialblast engages in transit amplifying mitotic divisions to produce increasing numbers of early stage germline cells, called spermatogonia. The cysts with the gonialblasts and the spermatogonia are located basal to the stem cells within the apical region of the testes. After ceasing amplifying divisions, the germline cells become spermatocytes, grow in size, and reduce their chromosomal content by meiosis. Once haploid, the germline cells compact their DNA and change their morphology from round to elongated cells, thereby becoming spermatids (Figure 1B). Fully developed sperm is found at the basal end of the testes (Fuller, 1993).

The two cyst cells surrounding each cluster of developing germline cells differentiate as well, as evident by their cellular growth and the expression of different sets of molecular markers (Zoller et al., 2012). CySCs express the transcription factor Zinc Finger Homeodomain-1 (Zfh-1) (Leatherman and DiNardo, 2008). The expression level of Zfh-1 drops in the CySC daughters as they differentiate and become displaced away from the stem cell area (Figure 1B, 1B’). CySCs also express the transcription factor Traffic Jam (Tj). Tj expression is maintained in cyst cells surrounding gonialblasts and spermatogonia (Figure 1B, 1B’) but drops in expression level in cyst cells that enclose spermatocytes (Li et al., 2003). Another transcription factor, Eyes absent (Eya), is expressed at low levels in CySCs, at higher levels in cyst cells surrounding spermatogonia, and at highest levels in cyst cells surrounding spermatocytes (Figure 1B, 1B’) (Fabrizio et al., 2003). The apical region of the testes in which Tj expression is high and Eya expression is increasing within the cyst cells coincides with the positions in which spermatogonia have reached the final two rounds of transit amplifying divisions and develop into spermatocytes. This area is, in the following, referred to as the transition zone (bracket in Figure 1B). Here, we show that Notch is expressed and activated in the cyst cells of the transition zone. We present evidence that the ligand Dl is required within the germline cells and that the receptor Notch is required within the cyst cells for the survival of the germline stem cell lineage.

## Results and Discussion

### The Notch pathway is activated in the transition zone of wild-type testes

Because germline cells and cyst cells are physically tightly associated, we reasoned that Notch signaling could play a role in germline-soma interactions. To explore this possibility, we first investigated the expression pattern of Notch in *wild-type* (*wt*) testes. For this, we employed two commonly used Notch antibodies: one directed against the NICD and the other against the Notch extra-cellular domain (NECD). Both antibodies were detected in the cyst cells surrounding the germline cells. To confirm expression in the cyst cells, we co-labeled the testes with the marker anti-Armadillo (Arm) that has previously been shown to localize to the membranes of the cyst cells, resulting in a net-like pattern of expression (Sarkar et al., 2007; Schulz et al., 2002). NICD (Figure 1C) and NECD (data not shown) were detectable in a similar net-like pattern both in the apical region where the cyst cells surround the spermatogonia (Figure 1C, arrows) and more basally where cyst cells surround the spermatocytes (Figure 1C, arrowheads). This expression pattern overlapped with the pattern produced by the Arm staining (Figure 1C’, 1C’’). We were not able to detect the expression of Dl by immuno-fluorescence staining of testes. However, western blots of testes revealed a band at the predicted size of about 62 kDa (data not shown), confirming its expression in the male gonad.

To confirm that Notch is active within the cyst cells and to address at which stage of cyst development Notch is activated, we used readouts for Notch stimulation. A Notch reporter, Notch Response Element-e-Green Fluorescent Protein (NRE-eGFP), consists of the GFP coding region under the control of Su(H) binding sites. Without Notch signaling, no GFP is expressed from the NRE-eGFP. When Notch is stimulated, Su(H), in combination with NICD, acts as an activator and GFP expression from NRE-eGFP is apparent (Zacharioudaki and Bray, 2014). Su(H)-driven GFP was not detectable above background in the cyst cells near the apical tip (Figure 1D, arrowhead). Instead, we detected strong GFP-signal in cyst cells starting more basally within the apical region (Figure 1D, arrows) and this expression was maintained in cyst cells throughout the testes (data not shown). Co-localization experiments revealed that NRE-eGFP is detectable in cyst cells of the apical region that also express Tj and Eya (Figure 1D’’ to 1D’’’, arrows). Though we cannot exclude the possibility that the expression of the NRE-eGFP at the tip of the testis is below detection level, our observations suggest that Notch most likely becomes first activated in the cyst cells of the transition zone (bracket in Figure 1B and 1D-D’’’).

### Over-activation of Notch in the cyst cells of Epidermal growth factor (EGF) mutant testes had a drastic effect on germline development

The Notch signaling pathway has been implicated in the specification of the male germline stem cell niche in *Drosophil*a males (Kitadate and Kobayashi, 2010). However, a role for Notch in the adult testes remained elusive, most likely because viable, temperature-sensitive alleles of Notch and many of the other tools (Table 1) that can be used to study Notch signaling in other tissues did not display a testis phenotype. This suggests that either loss of Notch has no effect on testes, or that it is extremely hard to eliminate Notch signaling from testes. Based on the above expression study, it appears that the area in which we first detect Notch activation is the same region of the testes in which EGF signaling is active. The EGF signaling pathway plays a major role in germline-soma interactions. First, signaling via EGF from the germline cells to the EGF-receptor (EGFR) on the cyst cells instructs cyst formation (Sarkar et al., 2007; Schulz et al., 2002). Subsequently, EGF signaling regulates cyst development. While a low dose of EGF from the germline to the cyst cells is required for the germline cells to progress through spermatogonial stages, a high dose of EGF signaling appears to promote the transition from early stage cysts containing spermatogonia into late stage cysts containing spermatocytes (Hudson et al., 2013, Kiger et al., 2000). A temperature-sensitive allele of EGF, *spitz*^*77-20*^, has served as an excellent tool for gaining more insight into spermatogenesis. Previous research has established that the *spitz*^*77-20*^ mutant testis phenotype from animals held at an intermediate temperature of 26.5°C could easily be modified by additional mutations in pathways regulating cyst development (Qian et al., 2014; Sarkar et al., 2007). Thus, prior to employing further tools for eliminating Notch signaling from otherwise *wt* testes we first utilized the *spitz*^*77-20*^ allele to investigate for a potential genetic interaction.

**Table 1:** Tools for studying Notch signaling. Fly stocks used to study Notch signaling in adult testes and their description, as indicated.

| BL# | Description | Genotype | Phenotype Observed |
| --- | --- | --- | --- |
| 29625 | UAS-*Cut*-RNAi | *y[1] v[1]; P{y[+t7.7] v[+t1.8]=TRiP.JF03304}attP2* | No |
| 33967 | UAS-*Cut*-RNAi | *y[1] sc[\*] v[1]; P{y[+t7.7]* | No |
| | | $v[+t1.8]=TRiP.HMS00924\}attP2$ | |
| 8611 | UAS-*Delta* with GFP inserted in extracellular domain | $w[*]; P\{w[+mC]=UAS-Dl::GFP\}DA55$ | No |
| 9319 | UAS-*Delta* | $y1 w^*; P\{UAS-dl.H\}2$ | No |
| 26697 | UAS-dominant negative *Delta* | $w^*; P\{UAS-Dl.DN\}TJ2/CyO$ | No |
| 26698 | UAS-dominant negative *Delta* | $w^*; P\{UAS-Dl.DN\}TJ3$ | No |
| 28032 | UAS-*Delta*-RNAi | $y1 v1; P\{TRiP.JF02867\}attP2$ | No |
| 34322 | UAS-*Delta*-RNAi | $y1 sc^* v1; P\{TRiP.HMS01309\}attP2$ | Yes |
| 36784 | UAS-*Delta*-RNAi | $y1 sc^* v1; P\{TRiP.GL00520\}attP40$ | No |
| 26322 | UAS-*Enhancer of split-m8*-RNAi | $y[1] v[1]; P\{y[+t7.7] v[+t1.8]=TRiP.JF02096\}attP2$ | No |
| 26675 | UAS-*Enhancer of split-mbeta* | $y[1] w[*]; P\{w[+mC]=UAS-E(spl)mbeta\}48.1$ | No |
| 26679 | UAS-*Enhancer of split-m4* | $w[*]; P\{w[+mC]=UAS-E(spl)m4-BFM.A\}15.5$ | No |
| 26872 | UAS-*Enhancer of split* | $w^*; P\{UAS-E(spl)m8-HLH.T\}3$ | No |
| 27179 | UAS-*E(spl)-malpha* | $y[1] w[*]; P\{w[+mC]=UAS-E(spl)malpha-BFM.A\}A17.4$ | No |
| 28735 | UAS-*Hintsight*-RNAi | $y[1] v[1]; P\{y[+t7.7] v[+t1.8]=TRiP.JF03162\}attP2$ | No |
| 33901 | UAS-*Hintsight*-RNAi | $y[1] sc[*] v[1]; P\{y[+t7.7] v[+t1.8]=TRiP.HMS00843\}attP2$ | No |
| 33943 | UAS-*Hintsight*-RNAi | $y[1] sc[*] v[1]; P\{y[+t7.7] v[+t1.8]=TRiP.HMS00894\}attP2/TM3, Sb[1]$ | No |
| 6578 | UAS-Dominant Negative *Kuzbanian* | $w[*]; P\{w[+mC]=UAS-kuz.DN\}2$ | No |
| 5830 | UAS-constitutively active *Notch* | $y1 w^*; P\{UAS-Dl::N.\Delta ECN\}B2a2$ | Yes |
| m7077 | UAS-*Notch* RNAi | $w^*; P\{UAS-N.dsRNA.P\}9G$ | No |
| 7078 | UAS-*Notch* RNAi | $P\{UAS-N.dsRNA.P\}14E, w^*$ | No |
| 26820 | UAS-*Notch* | $w^*; P\{UAS-Nfull\}3$ | No |
| 29856 | UAS-*Notch* truncated, ligand non-responsive | $P\{w[+mC]=UAS-N.DeltaEGF.LV\}1, w[1118]$ | No |
| 27988 | UAS-*Notch*-RNAi | $y[1] v[1]; P\{y[+t7.7] v[+t1.8]=TRiP.JF02959\}attP2$ | No |
| 28981 | UAS-*Notch*-RNAi | $y[1] v[1]; P\{y[+t7.7] v[+t1.8]=TRiP.JF01637\}attP2$ | No |
| 31180 | UAS-Notch-RNAi | $y[1] \ v[1]; P\{y[+t7.7] \ v[+t1.8]=TRiP.JF01693\}attP2$ | No |
| 31502 | UAS-Notch-RNAi | $y[1] \ v[1]; P\{y[+t7.7] \ v[+t1.8]=TRiP.JF01043\}attP2$ | No |
| 31503 | UAS-Notch-RNAi | $y[1] \ v[1]; P\{y[+t7.7] \ v[+t1.8]=TRiP.JF01053\}attP2$ | No |
| 33616 | UAS-Notch-RNAi | $\underline{y}1 \ v1; P\{TRiP.HMS00009\}attP2$ | No |
| 35640 | UAS-Notch-RNAi | $y[1] \ sc[*] \ v[1]; P\{y[+t7.7] \ v[+t1.8]=TRiP.GLV21004\}attP2$ | No |
| 36784 | UAS-Notch-RNAi | $y[1] \ sc[*] \ v[1]; P\{y[+t7.7] \ v[+t1.8]=TRiP.GL00520\}attP40$ | No |
| 31182 | UAS-Numb-RNAi | $y[1] \ v[1]; P\{y[+t7.7] \ v[+t1.8]=TRiP.JF01695\}attP2$ | No |
| 35045 | UAS-Numb-RNAi | $y[1] \ sc[*] \ v[1]; P\{y[+t7.7] \ v[+t1.8]=TRiP.HMS01459\}attP2$ | No |
| 28046 | UAS-mastermind-RNAi | $y[1] \ v[1]; P\{y[+t7.7] \ v[+t1.8]=TRiP.JF02881\}attP2$ | No |
| 26023 | UAS-neuralized-RNAi | $y[1] \ v[1]; P\{y[+t7.7] \ v[+t1.8]=TRiP.JF02048\}attP2$ | No |
| 35412 | UAS-neuralized-RNAi | $y[1] \ sc[*] \ v[1]; P\{y[+t7.7] \ v[+t1.8]=TRiP.GL00334\}attP2$ | No |
| 27498 | UAS-Nicastrin-RNAi | $y[1] \ v[1]; P\{y[+t7.7] \ v[+t1.8]=TRiP.JF02648\}attP2$ | No |
| 27681 | UAS-Presenilin-RNAi | $y[1] \ v[1]; P\{y[+t7.7] \ v[+t1.8]=TRiP.JF02760\}attP2/TM3, Sb[1]$ | No |
| 5815 | UAS-Serrate | $w[*]; P\{w[+mC]=UAS-Ser.mg5603\}SS1$ | No |
| 28713 | UAS-Serrate-RNAi | $y[1] \ v[1]; P\{y[+t7.7] \ v[+t1.8]=TRiP.JF03140\}attP2$ | No |
| 34700 | UAS-Serrate-RNAi | $y[1] \ sc[*] \ v[1]; P\{y[+t7.7] \ v[+t1.8]=TRiP.HMS01179\}attP2$ | No |
| 28900 | UAS-Su(H)-RNAi | $y[1] \ v[1]; P\{y[+t7.7] \ v[+t1.8]=TRiP.HM05110\}attP2/TM3, Sb[1]$ | No |
| 30727 | Expresses eGFP under control of NRE | $w[1118]; P\{w[+m*]=NRE-EGFP.S\}5A$ | N/A |
| 30728 | Expresses eGFP under control of NRE | $w[1118]; P\{w[+m*]=NRE-EGFP.S\}1$ | N/A |

In a *wt* testis, the germline cells vary in sizes and shapes and are easily recognizable by these characteristics. The GSCs and their transit amplifying daughters are small cells located within the apical region of the testis (Figure 2A, arrowhead), based on staining with the germline marker anti-Vasa. The spermatocytes are larger and found basal to the apical region and along the testis coil (Figure 2A, small arrows). Sperm is found at the testis base but also reaches into the lumen of the testis (Figure 2A, large arrow). We previously showed that when *spitz*^*77-20*^ mutant animals are raised at 26.5°C, the majority of the testes are tiny and contain mostly germline cells at the spermatogonia stage. These testes were classified as type I testes. Some testes are longer and also contain spermatocytes (type II testes) and/or spermatids (type III testes) (Qian et al., 2014). As most germline cells in *spitz*^*77-20*^ mutant testes are not enclosed by cyst cells, signaling between these two cell types via Notch and its transmembrane ligand seems unlikely. With this rationale, we employed the UAS/Gal4-expression system to express a ligand-independent and constitutively active version of Notch, UAS-*Dl::N.*Δ*ECN* (*caN*), within the cyst cells of *spitz*^*77-20*^ mutant testes (Brand and Perrimon, 1993; Duffy, 2002). This version of Notch contains the yeast Upstream Activating Sequences (UAS) as the promotor upstream of a fusion between the *dl* start and membrane transport signal sequence, and the coding region of the Notch transmembrane and intracellular domains (Baker and Schubiger, 1996).

**Fig. 2:**
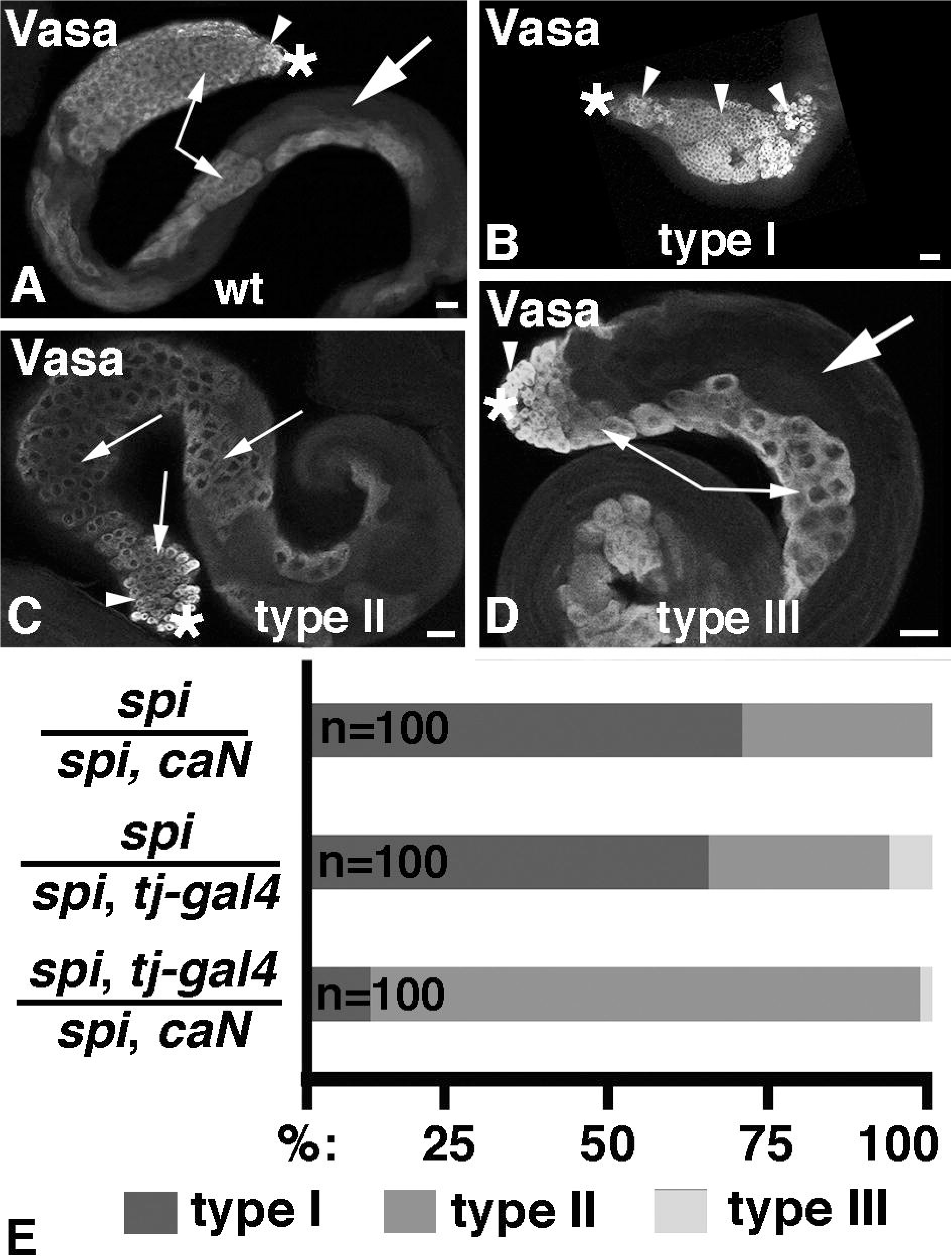
Activation of Notch within the cyst cells of EGF mutant testes modified the phenotype. A-D) Whole testes stained for the germline marker, Vasa. Asterisks mark the apical tips of the testes, arrowheads point to spermatogonia, small arrow point to spermatocytes, large arrows point to spermatids, scale bars: 30 μm. A) *wt*, B) *spi* type I, C) *spi* type II, and D) *spi* type III testis. E) Plot demonstrating the percentage of type I, II, and III testes in different genetic backgrounds (as indicated).

To express *caN* within the cyst cells of the testes, we used a well-established soma-Gal4 transactivator, *traffic jam-gal4* (*tj-gal4*) (Hayashi et al., 2002). As expected, control animals (*spi/spi, caN* and *spi/spi, tj-gal4*) had mostly type I testes (Figure 2E, n=100) that were filled with spermatogonia (arrowheads in Figure 2B). Upon expression of *caN* in the cyst cells of *spitz*^*77-20*^ animals, we detected a drastic modification of the mutant phenotype. Almost all of the experimental testes (*spi, tj-gal4/spi, caN*, n=100) were type II testes (Figure 2E) and contained mostly spermatocytes (small arrows in Figure 2C). A few testes were of type III (Figure 2E) and contained sperm bundles (large arrow in Figure 2D). We do not understand how expression of *caN* allows the cells in *spitz*^*77-20*^ testes to develop past the spermatogonial stage. However, this finding suggests that Notch does play a role in cyst development, likely by promoting germline differentiation, and encouraged us to further explore the tools for generating a Notch loss-of-function phenotype.

### Reduction of Dl from the germline caused massive loss of germline cells

The expression pattern of Notch and Notch reporters suggests that the cyst cells receive the ligand from the germline. As *dl* mutation are embryonic lethal, we used the FRT/Flp-recombination technique to generate negatively GFP-marked clusters of *dl* mutant germline cells in adult testes (Xu and Rubin, 1993). In the control animals, 18 out of 46 testes contained one or more GFP-negative clusters of germline cells. In the experimental animals, only three out of 80 testes showed a single cluster of GFP-negative germline cells (data not shown). This suggests that *dl* mutant germline clones are either rarely formed or rarely survive.

To obtain higher numbers of *dl* mutant cells, we used a RNA-Interference (RNAi) construct for *dl* (*UAS-dl-i*) driven either in cyst cells via *tj-gal4*, or within the germline via the *nanos-gal4* transactivator (*nos-gal4*; Van Doren et al., 1998). While expression of UAS-*dl-i* via *tj-gal4* had no morphological effect, expression of UAS-*dl-i* via *nos-gal4* caused a drastic loss of germline cells. When *nos-gal4/UAS-dl-i* animals were kept at the permissive temperature of 18°C, no effect on the testes was observed (n=30, data not shown). After shifting the animals to the restrictive temperature of 29°C for eight days, testes were long and thin and contained only few germline cells (Figure 3A, n=50) while testes from control animals (*nos-gal4/wt, n=30* and *UAS-dl-i/wt*, n=30) treated under the same conditions appeared normal and contained all stages of development (Figure 3B). Specifically, GSCs (arrowhead in Figure 3B), spermatogonia (small arrows in Figure 3B) and spermatocytes (large arrow in Figure 3B) filled the apical region of the testes from *nos-gal4/wt* animals.

**Fig. 3:**
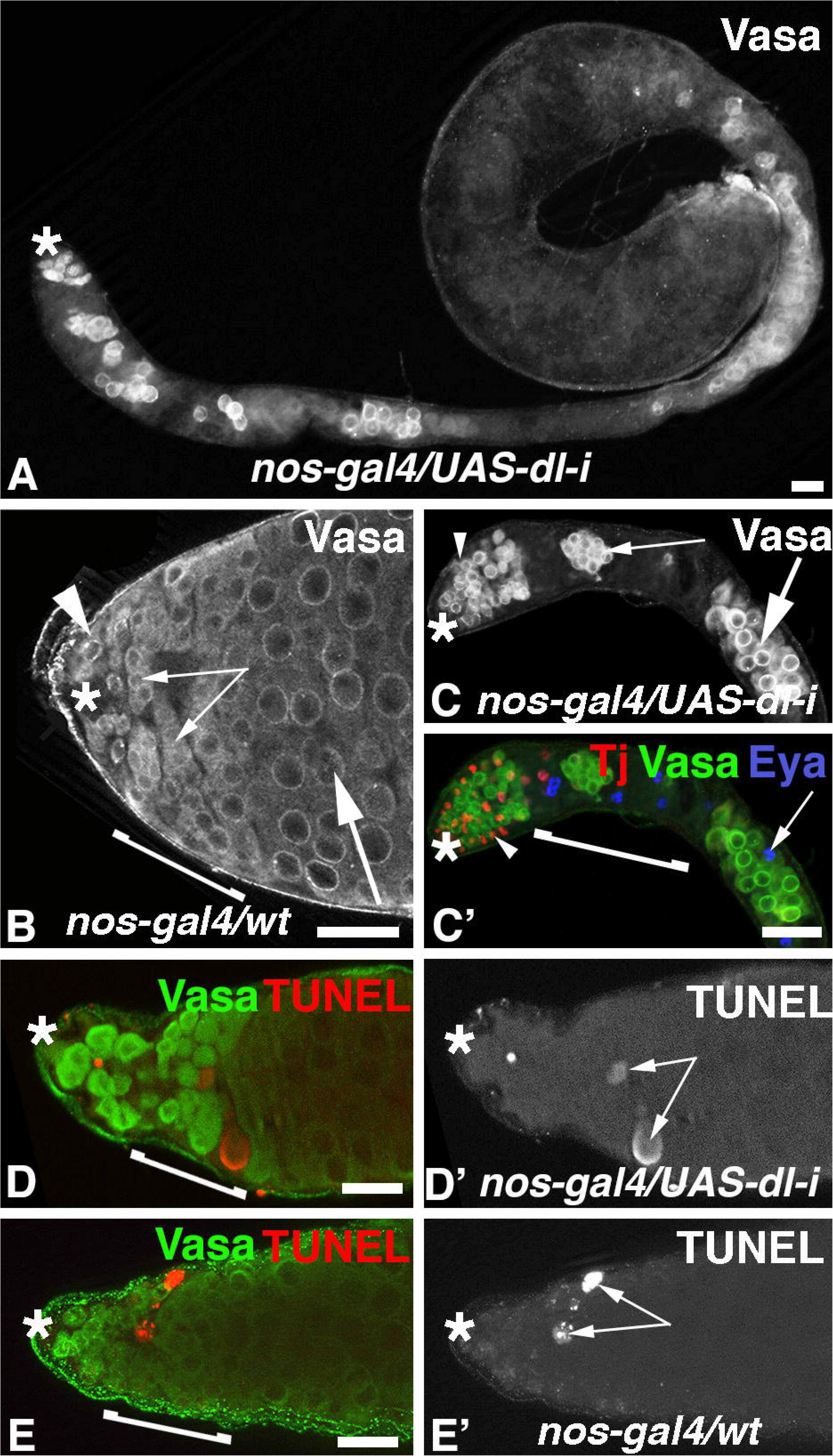
Knockdown of *dl* in the germline caused severe germline loss. A-E) Scale bars: 30 μm, asterisks mark the apical tips of the testes, bracket: transition zone, stainings as indicated. A) A testis with knockdown of *dl* in the germline after shifting males to the restrictive temperature for eight days. B, C) The apical region of B) a control testis and C) a testis with knockdown of *dl* in the germline. Arrowheads point to GSCs and/or their immediate daughters, small arrows point to spermatogonia, large arrows point to spermatocytes. C’) Same apical testis region as in C) but co-labeled for cyst cell markers. Arrowhead points to a Tj-positive cyst cell nucleus, arrow points to an Eya-positive cyst cell nucleus. D-D’) The apical region of a testis with *dl* knock-down in the germline. E-E’) The apical region of a control testis.

By eight days after the temperature shift to 29°C, the testes from *nos-gal4/UAS-dl-i* animals had GSCs and gonialblasts next to the hub (Figure 3C, arrowhead, n=45), but contained only a few clusters of spermatogonia (Figure 3C, small arrow) and/or spermatocytes (Figure 3C, large arrow). The somatic cells, the hub cells and the cyst cells, were present in testes from *nos-gal4/UAS-dl-i* animals. In the apical region, cyst cells expressing Tj (Figure 3C’, arrowhead) were intermingled with the germline cells. Cyst cells expressing Eya were also detected and some of them appeared to be associated with germline cells (Figure 3C’, arrow). This suggests that the germline was not lost due to the lack of somatic cells.

To determine how the germline was lost we performed a time-line experiment by shifting *nos-gal4/UAS-dl-i* animals to the restrictive temperature for three to eight days. Testes were labeled with the germline marker, anti-Vasa, and the cell death marker, TUNEL, at each day of the experiment. By five days after the temperature shift, testes from *nos-gal4/UAS-dl-i* animals had fewer Vasa-positive cells within the transition zone (Figure 3D, bracket) and showed many TUNEL-positive spots in this area instead (Figure 3D’, arrows). As we also detected TUNEL-positive spots within the transition zone of control testes from *nos-gal4/wt* animals (Figure 3E, 3E’), we compared the number of TUNEL-positive spots in the transition zone of testes from *nos-gal4/UAS-dl-i* and *nos-gal4/wt* animals. A detailed analysis revealed significantly increased numbers of TUNEL-positive spots in testes from *nos-gal4/UAS-dl-i* animals compared to the control testes from *nos-gal4/wt* animals starting at five days after the temperature shift (Figure 4A). We conclude that *dl* acts within the germline cells for their survival.

**Fig. 4:**
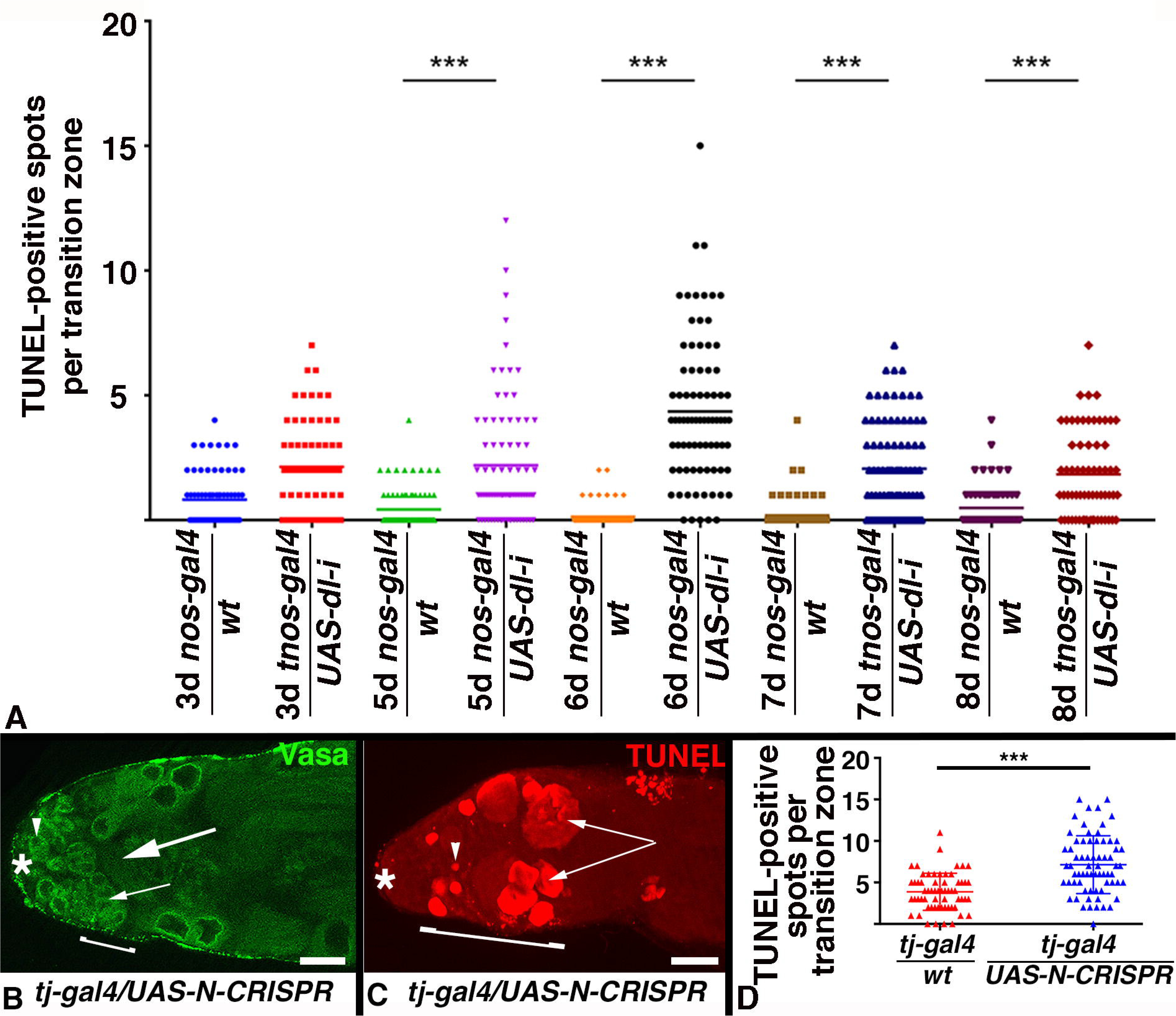
Loss of Notch signaling caused massive cell death. A) Plot showing the numbers of TUNEL-positive spots within the transition zone of control and *dl* knockdown animals at various time points after the shift to the restrictive temperature. Asterisks indicate statistical significance, P-value <0.001. B, C) The apical regions of testes from animals expressing UAS-N-CRISPR within the cyst cells, asterisks mark the tips of the testes, scale bars: 30 μm, brackets: transition zones, staining as indicated. B) Note that the mutant testis contains only a few spermatogonia (small arrows). Arrowhead points to a GSC, large arrow points to a sperm bundle in the apical regions. C) The mutant testis shows massive cell death. Arrowhead points to a single TUNEL-positive cell, arrows point to clusters of TUNEL-positive cells. D) Plot showing the numbers of TUNEL-positive spots within the transition zone of control and UAS-N-CRISPR animals at two weeks after the shift to the restrictive temperature. Asterisks indicate statistical significance, P-value <0.001.

### Loss of Notch from the soma caused cell death

The *Notch* gene maps to the X-chromosome, making it impossible to use a simple FRT/Flp-technique for the generation of mutant clones, and a more complex system is currently not available. Furthermore, none of the viable *Notch* mutant alleles nor the expression of *Notch*-RNA*i*-constructs produced a phenotype in testes (Table 1). Likewise, modulating the expression of signal transducers known to act downstream of Notch via RNA*i* did not produce a phenotype in our hands (Table 1). Therefore, we employed the CRISPR technology in combination with the UAS/Gal4-system (Gratz et al., 2015). A UAS-Notch-CRISPR line was previously reported to efficiently reduce Notch expression and cause the expected mutant phenotype in the wing discs (Gao et al., 2015). Expressing this UAS-Notch-CRISPR in the cyst cells of the testes (*tj-gal4/UAS-N-CRISPR)* for 14 days at 29°C produced a similar phenotype as seen in testes from *nos-gal4/UAS-dl-i* animals revealing that Notch acts within the cyst cells. Based on the expression of Vasa, testes from *tj-gal4/UAS-N-CRISPR* animals lost the germline within the transition zone (Figure 4B), while testes from control animals did not (data not shown). TUNEL analysis at 14 days after temperature shift revealed excessive cell death in the apical region of the testes from *tj-gal4/UAS-N-CRISPR* animals (Figure 4C) and the number of TUNEL-positive spots significantly exceeded the number of spots in control testes (*tj-Gal4/wt*, Figure 4D). The reduction in germline cells and the increase in TUNEL-positive spots in testes from *nos-gal4/UAS-dl-i* and *tj-gal4/UAS-N-CRISPR* animals coincides with the expression of the NRE-eGFP reporter in the transition zone and suggests that Notch signaling is essential for the survival of germline cells that have left the stem cell area and are transitioning towards differentiation. Our findings add yet another example to the literature where Notch signaling can have different effects within the same tissue. During testis development, Notch signaling is essential for the fate specification of the stem cell niche, but in the adult testis Notch signaling is required for germline survival.

Though our data demonstrate that Notch signaling is required for germline survival, the molecular mechanism underlying this effect remains elusive. It is well established that Notch receptor-ligand interaction provides a strong adhesive force between two communicating cells (Murata and Hayashi, 2016). Failure to maintain cell adhesion may cause or contribute to the inability of the cysts to differentiate in unison and result in death when Notch signaling is lost. The idea that Notch and Dl act via cell adhesion is consistent with our failure to detect a loss-of-function phenotype by reducing any of the other signaling molecules of the canonical pathway (Table 1). However, it is possible that the available tools are not efficient in testes. Alternatively, Notch could act in combination with a different set of molecules in the testes. In the nervous system, a non-canonical role for Notch has been demonstrated. Notch genetically and physically interacts with Disabled and Trio, both of which are components of Abelson tyrosine kinase (Abl) signaling pathway (LeGall et al., 2008). Abl is a non-receptor tyrosine kinase that has been implicated in cell contact, morphogenesis, growth, and migration (Bradley and Koleske, 2009). Specifically, Abl promotes cell adhesion via cadherin-based cell contacts (Zandy and Pendergast, 2008; Zandy et al., 2007). Thus, it is possible that Notch acts in a similar manner in the cysts of the testes to maintain the intimate contact between germline cells and surrounding cyst cells.

## Acknowledgements

We thank Guanjun Gao, Dorothea Godt, and Zhaoyu Xue for flies and reagents, Alicia Hudson and Leon McSwain for technical assistance, and Manashree Malpe and Karl Kudyba for critical reading of the manuscript. This work was supported by NSF grants #0841419 and #1355099 given to CS.

## Materials and Methods

### Fly husbandry

Flies were maintained on standard cornmeal/molasses diet at room temperature. *tj-gal4* was obtained from the Kyoto stock center and all other stocks from the Bloomington *Drosophila* Stock Center (BDSC; genotypes and stock numbers are listed in Table 1).

### Immunofluorescence and image analysis

Testes were isolated according to Parrott et al. (2012). The mouse anti-Eyes absent (Eya) antibody (1:10) developed by S. Benzer and N. M. Bonini, and the rat anti-Vasa antibody (1:10) developed by A. C. Spradling and D. Williams were obtained from the Developmental Studies Hybridoma Bank, created by the NICHD of the NIH and maintained at The University of Iowa, Department of Biology, Iowa City, IA 52242. Goat anti-Armadillo antibody (1:200) was obtained from Santa Cruz Biotechnology Inc. (sc28653), and the rabbit anti-Green Fluorescent Protein (GFP) antibody (1:200) was obtained from Life Technologies (A11122), Guinea pig-anti-Traffic Jam antibody (1:5000) was a gift from Dorothea Godt. Secondary Alexa 488, 568, and 647-coupled antibodies (1:1000) and Slow Fade Gold embedding medium were obtained from Life Technologies. Images were taken using a Zeiss Axiophot with a digital camera and apotome and processed using Axiovision Rel. software. Images were analyzed using ImageJ and processed with Photoshop. Statistical relevance was determined using the Graphpad student’s t-test.

### UAS/Gal4-expression studies and CRISPR

Fly crosses were set up at 18°C and progeny shifted as larvae or adults to the restrictive temperatures as described in the result and discussion section of the manuscript.

### Apoptosis detection

Cells in apoptosis were detected using terminal deoxynucleotidyl transferase dUTP nick end labeling (TUNEL). The Apoptag Red In Situ Kit was obtained from Millipore (S1765) and tissue was treated according to the manufacturer’s instructions.

## References

Baker, R. and Schubiger, G. (1996). Autonomous and nonautonomous Notch functions for embryonic muscle and epidermis development in Drosophila. Development 122, 617–626.

Berset, T., Hoier, E. F., Battu, G., Canevascini, S. and Hajnal, A. (2001). Notch inhibition of RAS signaling through MAP kinase phosphatase LIP-1 during C. elegans vulval development. Science 291, 1055–1058.

Bradley, W. D. and Koleske, A. J. (2009). Regulation of cell migration and morphogenesis by Abl-family kinases: emerging mechanisms and physiological contexts. J Cell Sci 122, 3441–3454.

Brand, A. H. and Perrimon, N. (1993). Targeted gene expression as a means of altering cell fates and generating dominant phenotypes. Development 118, 401–415.

Bray, S. J. (2016). Notch signalling in context. Nat Rev Mol Cell Biol 17, 722–735.

Dorsky, R. I., Chang, W. S., Rapaport, D. H. and Harris, W. A. (1997). Regulation of neuronal diversity in the Xenopus retina by Delta signalling. Nature 385, 67–70.

Duffy, J. B. (2002). GAL4 system in Drosophila: a fly geneticist’s Swiss army knife. Genesis 34, 1–15.

Fabrizio, J. J., Boyle, M. and DiNardo, S. (2003). A somatic role for eyes absent (eya) and sine oculis (so) in Drosophila spermatocyte development. Dev Biol 258, 117–128.

Fairchild, M. J., Islam, F. and Tanentzapf, G. (2017). Identification of genetic networks that act

Fuller, M. T. (1993). Spermatogenesis in *Drosophila*.. In The development of Drosophila melanogaster. (ed. M. Bate, Martinez Arias, A.), pp. 71–148. Cold Spring Harbor, New York, USA: Cold Spring Harbor Laboratory Press.

Gao, Y., Liu, T. and Huang, Y. (2015). MicroRNA-134 suppresses endometrial cancer stem cells by targeting POGLUT1 and Notch pathway proteins. FEBS Lett 589, 207–214.

Gratz, S. J., Rubinstein, C. D., Harrison, M. M., Wildonger, J. and O’Connor-Giles, K. M. (2015). CRISPR-Cas9 Genome Editing in Drosophila. Curr Protoc Mol Biol 111, 31 32 31–20.

Griswold, M. D. (1998). The central role of Sertoli cells in spermatogenesis. Semin Cell Dev Biol 9, 411–416.

Guruharsha, K. G., Kankel, M. W. and Artavanis-Tsakonas, S. (2012). The Notch signalling system: recent insights into the complexity of a conserved pathway. Nat Rev Genet 13, 654–666.

Hardy, R. W., Tokuyasu, K. T., Lindsley, D. L. and Garavito, M. (1979). The germinal proliferation center in the testis of Drosophila melanogaster. J Ultrastruct Res 69, 180–190.

Hayashi, S., Ito, K., Sado, Y., Taniguchi, M., Akimoto, A., Takeuchi, H., Aigaki, T., Matsuzaki, F., Nakagoshi, H., Tanimura, T., et al. (2002). GETDB, a database compiling expression patterns and molecular locations of a collection of Gal4 enhancer traps. Genesis 34, 58–61.

Hori, K., Sen, A. and Artavanis-Tsakonas, S. (2013). Notch signaling at a glance. J Cell Sci 126, 2135–2140.

Hudson, A. G., Parrott, B. B., Qian, Y. and Schulz, C. (2013). A temporal signature of epidermal growth factor signaling regulates the differentiation of germline cells in testes of Drosophila melanogaster. PLoS One 8, e70678.

Jain, R., Rentschler, S. and Epstein, J. A. (2010). Notch and cardiac outflow tract development. Ann N Y Acad Sci 1188, 184–190.

Kiger, A. A., White-Cooper, H. and Fuller, M. T. (2000). Somatic support cells restrict germline stem cell self-renewal and promote differentiation. Nature 407, 750–754.

Kitadate, Y. and Kobayashi, S. (2010). Notch and Egfr signaling act antagonistically to regulate germ-line stem cell niche formation in Drosophila male embryonic gonads. Proc Natl Acad Sci U S A 107, 14241–14246.

Kopan, R. and Ilagan, M. X. (2009). The canonical Notch signaling pathway: unfolding the activation mechanism. Cell 137, 216–233.

Le Gall, M., De Mattei, C. and Giniger, E. (2008). Molecular separation of two signaling pathways for the receptor, Notch. Dev Biol 313, 556–567.

Leatherman, J. L. and Dinardo, S. (2008). Zfh-1 controls somatic stem cell self-renewal in the Drosophila testis and nonautonomously influences germline stem cell self-renewal. Cell Stem Cell 3, 44–54.

Li, M. A., Alls, J. D., Avancini, R. M., Koo, K. and Godt, D. (2003). The large Maf factor Traffic Jam controls gonad morphogenesis in Drosophila. Nat Cell Biol 5, 994–1000.

MacGrogan, D., Nus, M. and de la Pompa, J. L. (2010). Notch signaling in cardiac development and disease. Curr Top Dev Biol 92, 333–365.

Mason, H. A., Rakowiecki, S. M., Gridley, T. and Fishell, G. (2006). Loss of notch activity in the developing central nervous system leads to increased cell death. Dev Neurosci 28, 49–57.

Matunis, E., Tran, J., Gonczy, P., Caldwell, K. and DiNardo, S. (1997). punt and schnurri regulate a somatically derived signal that restricts proliferation of committed progenitors in the germline. Development 124, 4383–4391.

Murata, A. and Hayashi, S. (2016). Notch-Mediated Cell Adhesion. Biology (Basel) 5.

Nefedova, Y., Sullivan, D. M., Bolick, S. C., Dalton, W. S. and Gabrilovich, D. I. (2008). Inhibition of Notch signaling induces apoptosis of myeloma cells and enhances sensitivity to chemotherapy. Blood 111, 2220–2229.

Parrott, B. B., Hudson, A., Brady, R. and Schulz, C. (2012). Control of germline stem cell division frequency--a novel, developmentally regulated role for epidermal growth factor signaling. PLoS One 7, e36460.

Qian, Y., Dominado, N., Zoller, R., Ng, C., Kudyba, K., Siddall, N. A., Hime, G. R. and Schulz, C. (2014). Ecdysone signaling opposes epidermal growth factor signaling in regulating cyst differentiation in the male gonad of Drosophila melanogaster. Dev Biol 394, 217–227.

Sarkar, A., Parikh, N., Hearn, S. A., Fuller, M. T., Tazuke, S. I. and Schulz, C. (2007). Antagonistic roles of Rac and Rho in organizing the germ cell microenvironment. Curr Biol 17, 1253–1258.

Schulz, C., Wood, C. G., Jones, D. L., Tazuke, S. I. and Fuller, M. T. (2002). Signaling from germ cells mediated by the rhomboid homolog stet organizes encapsulation by somatic support cells. Development 129, 4523–4534.

Tran, J., Brenner, T. J. and DiNardo, S. (2000). Somatic control over the germline stem cell lineage during Drosophila spermatogenesis. Nature 407, 754–757.

Van Doren, M., Williamson, A. L. and Lehmann, R. (1998). Regulation of zygotic gene expression in Drosophila primordial germ cells. Curr Biol 8, 243–246.

Wang, S., Sdrulla, A. D., diSibio, G., Bush, G., Nofziger, D., Hicks, C., Weinmaster, G. and Barres, B. A. (1998). Notch receptor activation inhibits oligodendrocyte differentiation. Neuron 21, 63–75.

Xu, T. and Rubin, G. M. (1993). Analysis of genetic mosaics in developing and adult Drosophila tissues. Development 117, 1223–1237.

Yamashita, Y. M., Jones, D. L. and Fuller, M. T. (2003). Orientation of asymmetric stem cell division by the APC tumor suppressor and centrosome. Science 301, 1547–1550.

Ye, Q. F., Zhang, Y. C., Peng, X. Q., Long, Z., Ming, Y. Z. and He, L. Y. (2012). Silencing Notch-1 induces apoptosis and increases the chemosensitivity of prostate cancer cells to docetaxel through Bcl-2 and Bax. Oncol Lett 3, 879–884.

Zacharioudaki, E. and Bray, S. J. (2014). Tools and methods for studying Notch signaling in Drosophila melanogaster. Methods 68, 173–182.

Zandy, N. L. and Pendergast, A. M. (2008). Abl tyrosine kinases modulate cadherin-dependent adhesion upstream and downstream of Rho family GTPases. Cell Cycle 7, 444–448.

Zandy, N. L., Playford, M. and Pendergast, A. M. (2007). Abl tyrosine kinases regulate cell-cell adhesion through Rho GTPases. Proc Natl Acad Sci U S A 104, 17686–17691.

Zoller, R. and Schulz, C. (2012). The Drosophila cyst stem cell lineage: Partners behind the scenes? Spermatogenesis 2, 145–157.

Zweidler-McKay, P. A., He, Y., Xu, L., Rodriguez, C. G., Karnell, F. G., Carpenter, A. C., Aster, J. C., Allman, D. and Pear, W. S. (2005). Notch signaling is a potent inducer of growth arrest and apoptosis in a wide range of B-cell malignancies. Blood 106, 3898–3906.

